# Specificity of *Drosophila innubila* Nudivirus Infection in *Drosophila* Cell Culture

**DOI:** 10.1101/2025.06.09.658616

**Authors:** Kent M. Mulkey, Taiye Adewumi, Robert L. Unckless

## Abstract

Viral host range is an important aspect of both viral biology but also the pragmatic issue of producing viral stocks for experimentation. Host range is important both in terms of the species a virus can infect (taxonomy), and the types of host cells the virus can infect (tropism). Nudiviruses are large DNA viruses that infect several arthropods and are generally poorly studied. Several nudiviruses infect *Drosophila* species including the *Drosophila innubila* Nudivirus (DiNV). We aimed to identify cell lines that support the replication of DiNV both for the sake of understanding host range, and to develop a system for large scale production of the virus for infection. We utilized cell lines from the focal host, *D. innubila*, as well as available cell lines from *D. virilis* and *D. melanogaster* and inoculated with both a wild-collected pool of “naïve” virus and a *D. innubila* cell culture-adapted isolate. We found that virus from wild caught flies infected cells from the 3 cell lines and replicate its genome, but the passage 1 fluids from these infections were unable to reinfect upon introduction to new cells. In contrast, a cell culture-adapted strain of DiNV infected the same *Drosophila* cells (though relatively poorly in *D. melanogaster* cells) and produced infectious progeny that infected new cells. Thus, our cell culture-adapted virus developed the ability to infect broadly *and* produce infectious virions.

## Introduction

Arthropods serve as excellent models to study host/pathogen interaction since they are genetically tractable, inexpensive to rear and have short lifespans that allow longitudinal experiments (Sabin et al., 2010; Wang et al., 2010). *Drosophila* has long been at the forefront of this research (Hughes et al., 2012), but surprisingly, DNA viruses were historically absent. Several RNA viruses are known to naturally infect *Drosophila* and many of these are well studied (David M. Knipe, 2013; Tafesh-Edwards and Eleftherianos, 2020). Most lab studies with DNA viruses, however, have utilized viruses that do not naturally infect *Drosophila* (Webster et al., 2015; Xu and Cherry, 2014). In 2011 a DNA virus from the Nudivirus family, *Drosophila innubila* Nudivirus (DiNV), was discovered to naturally infect *Drosophila innubila* which inhabits in the Sky Islands in the Southwestern United States (Unckless, 2011). A few years later, *D. melanogaster* was found to host Kallithea virus, a close relative of DiNV, and more recently several more DNA viruses of *Drosophila* were discovered (Wallace et al., 2021; Webster et al., 2015). To date, however, researchers have lacked a good cell culture system for studying nudiviruses that infect *Drosophila*.

Nudiviruses are a family of enveloped, rod shaped, double stranded DNA viruses that infect a diversity of terrestrial and aquatic arthropods (Couch, 1974; Holt et al., 2019; Huger, 1966; Petersen et al., 2022; Raina, 2000; Unckless, 2011). DiNV is typical of nudiviruses and has a circular double-stranded DNA genome of about 155kb and encodes for 107 open reading frames. DiNV’s genome shares about 35 core genes with other Nudiviruses and 21 orthologous genes with the insect infecting Baculoviruses (Petersen et al., 2022). DiNV is closely related to the better-studied *Oryctes rhinoceros* Nudivirus (OrNV) which has been utilized for biological control of the coconut palm rhinoceros beetle, *Oryctes rhinoceros* (Huger, 1966; Payne, 1974). Originally, because there was not a cell culture system available, the virus used in the biocontrol was cultivated in live beetle larvae. Then, in 1982 a cell line derived from *Heteronychus arator* (HA1179), a Coleopteran Scarab beetle, was described (Crawford, 1982) and reported to support OrNV replication (Crawford and Sheehan, 1985). Three isolates of OrNV were grown in the HA1179 cells to titers between 10^5^ and 10^7^ ID_50_/ml (Khudri et al., 2016). The ability to grow OrNV in cell culture has benefited the biocontrol effort with a virus production method that is more economical and controlled than virus production in the live insect larvae.

Due to our inability to culture DiNV in the lab, most of our understanding to date comes from studies in the wild of both life history traits and genome sequence diversity (Hill and Unckless, 2018, 2020; Unckless, 2011). Overall, the virus is both common, consistently found in more than 30% of wild-caught flies (by PCR screening) and virulent, both killing and reducing fecundity of infected flies (Hill and Unckless, 2020; Unckless, 2011). High virulence appears to evolve repeatedly through a series of mutations across the viral genome. The closely related Kallithea virus encodes a suppressor of host Toll signaling which likely contributes to virulence (Palmer et al., 2019; Palmer et al., 2018). In the wild, the high virulence haplotype of DiNV is associated with lower expression of Toll-related genes in the host (Hill and Unckless, 2020).

Being able to grow viruses in the lab with a cell culture system provides a source of virus that contains minimal extraneous or contaminating organisms (Leland and Ginocchio, 2007; Lynn, 1999). We evaluated three Drosophila cell lines for their ability to grow DiNV. We inoculated cell lines with a naïve wild-isolated DiNV or a cell culture-adapted isolate of DiNV and assayed for DiNV genome increase after 10 days of incubation using qPCR (Watzinger et al., 2006). For each virus isolate we determined the extent to which we could detect viral genome increase 10 days after an initial inoculation of the cells (Passage 1) and after a second inoculation of clean cells with passage 1 fluids (Passage 2). Overall, we found that while the naïve virus could replicate its genome, it could not be passaged. In contrast, the cell-culture adapted virus both replicated its genome and was able to be passaged – particularly in the native *D. innubila* cells and the relatively closely related *D. virilis* cells.

## Materials and Methods

### Cells

The cells that were used for these studies included Dv-1 (*Drosophila virilis*, obtained from the Drosophila Genomics Resource Center, Bloomington, IN), S2 (*Drosophila melanogaster*, obtained from Kausik Si, Stowers Research Institute, Kansas City, MO) and Dinn-1 (*Drosophila innubila*, (Schedl et al., 2025)).

All cells were maintained in Schneiders Drosophila Medium (Gibco) supplemented with heat inactivated fetal bovine serum (Gibco or Sigma, 10% final concentration) and gentamicin (Gibco, 10 ng/ml). Dv-1 cells were passaged once a week with a 1:20 split ratio, S2 cells were passaged once a week with a 1:10 split ratio, and Dinn-1 cells were passaged once every 2 weeks with a 1:3 split ratio. The Dinn-1 cells require trypsin treatment for passage. For routine culture, all three cell types were grown in Corning or Falcon 25 cm^2^ tissue culture flasks with a plug seal cap and maintained at 23°C in a non-humidified incubator. For simplicity, we refer to these cells throughout as Di (Dinn-1 from *D. innubila*), Dv (Dv-1 from *D. virilis*) and Dm (S2 from *D. melanogaster*).

### Viral isolates

The experiments used virus derived immediately from fly homogenate or from cell-culture adapted virus (adapted from the fly homogenate). The fly homogenate was prepared as follows: individual flies collected from the Chiricahua mountains of Arizona in September 2018 were placed in a microcentrifuge tube. One hundred microliters of viral buffer (Unckless, 2011) was added to each tube and the flies were homogenized with a handheld tissue pestle. DNA was extracted from 50 μL of the homogenate screened for DiNV by PCR as described in Hill and Unckless (2020). The virus positive fluids were then pooled and brought up to 5 ml in Schneiders medium with 10 ng/mL gentamicin and then passed through a 0.45μM membrane filter to remove cellular debris and bacteria. The virus fluids were aliquoted to 200 μL and stored at-80°C. Fluids isolated after a single passage were used for a passage 2 experiment (see below). Cell culture adapted-virus was isolated after a sample of progenitor Dinn-1 cells were inoculated with the fly homogenate fluids and the culture incubated for about one year with routine fluid changes (Schedl et al., 2025). Passage 2 from the cell culture-adapted virus was collected as described for the fly homogenate passage 2 virus.

### Growth Study Protocol

We conducted 4 growth studies. Each study involved inoculating the cells with virus and using qPCR to determine the viral genome increase after 10 days of incubation. We therefore use viral nucleic acid copy number as a proxy for viral titer. This was necessary because we have not observed cytopathic effect from DiNV, ruling out plaque assays, and at the time of the experiment, we had no way to count infectious particles for other count-based assays (but see below). The first study used virus derived from wild-caught fly homogenate (naïve virus) to inoculate the cells. The second study involved inoculating new cells with the passaged fluids generated in the first study. The third study used the cell culture derived isolate of the virus, and the fourth study used the passaged fluids from the third study. The first growth study was conducted in two complete blocks, generating 8 samples (4 from each block) for each set of treatment conditions which consisted of a 0-hour mock inoculated cell control group, a 240-hour mock inoculated cell control group, a 0-hour virus inoculated group, and a 240-hour virus inoculated group. The second, third and fourth studies were conducted in a single block with 4 samples generated for each treatment condition. For each experiment, cells were seeded into 48 well plates at a density of approximately 1 x 10^5^ to 5 x 10^5^ cells/well then 24 hours later 4 wells of each cell were inoculated with approximately 2400 fluorescent focus forming units for each treatment condition. We inoculated 2 identical plates for each condition with one being frozen (-80°C) immediately after addition of virus (0-hour sample) and the other was frozen after 240 hours of incubation (240-hour sample).

### Sample Preparation

Frozen samples were thawed at room temperature. The contents of each well were homogenized with a pipettor under sterile conditions and then a 50 μL sample was collected and added to a clean Eppendorf microcentrifuge tube. One microliter of diluted lambda DNA was then added to the 50 μL of sample to serve as an internal extraction control. DNA was extracted from prepared cell/virus lysates using the Qiagen Puregene Cell Core DNA Kit (Qiagen, Hilden, Germany) following manufacturer’s recommendations for cell culture.

### Quantitative PCR to quantify viral and host nucleic acids

The quantitative polymerase chain reaction (qPCR) was performed using the SsoAdvanced Universal SYBR detection system with the CFX Connect Real Time PCR System and the Bio-Rad CFX Manager 3.1 (all three are Bio-Rad, Hercules, CA, USA). Each reaction mixture contained 1 μL of extracted target DNA, 0.4 μM of each gene-specific forward and reverse primer in 0.5 μL water, 5 μL of 2x SYBR green Supermix and Molecular Biology grade water for a total reaction volume of 10 μl. The thermal cycling conditions were: 3 minutes at 95°C, followed by 32 cycles of 10 seconds at 95°C and 30 seconds at 60°C and then a final dissociation-curve step of 5 minutes at 95°C, 5 seconds at 60°C and 15 seconds at 95°C. All primers are listed in Table S1. To determine the qPCR efficiency of each primer pair, 2-fold or 10-fold serial dilutions of DNA were used as templates to construct the standard curve. The standard curve of each primer pair was generated based on the linear relationship between the mean threshold cycle (Cq) values and the dilutions. Calculations were performed using Microsoft Excel. The amplification efficiency was determined using the slopes of the standard curves (Svec et al., 2015).

### Verification of viral transcription in passage 2 experiments with RT-PCR

We utilized reverse-transcriptase PCR (RT-PCR) to confirm the presence of viral transcripts in samples. Cells from each treatment group (fly homogenate and cell culture–adapted virus) were harvested at two timepoints (0 hr and 240 hr post inoculation) from each cell line (Di, Dv, and Dm), and total RNA was extracted. Briefly, 500 µL of cold TRIzol reagent was added to each sample and homogenized. Samples were incubated at room temperature for 5 minutes, followed by the addition of 100 µL of cold chloroform and another 3-minute incubation at room temperature. Samples were then centrifuged at 12,000 × g for 15 minutes at 4 °C. The aqueous (top) phase was carefully transferred to new tubes. Glycogen and 100% isopropanol were added to each tube, followed by a 10-minute incubation at room temperature. Samples were centrifuged at 12,000 × g for 10 minutes at 4 °C. The supernatant was discarded, and the RNA pellet was washed with 75% ethanol by centrifugation at 7,500 × g for 5 minutes at 4 °C. After removing the ethanol, pellets were air-dried and resuspended in nuclease-free water. Samples were incubated at 55 °C for 10 minutes to facilitate resuspension.

Genomic DNA contamination was removed using the Invitrogen™ DNA-free™ DNA Removal Kit and RNA concentration was quantified using a Qubit fluorometer. Complementary DNA (cDNA) was synthesized from purified RNA using the iScript™ Reverse Transcription Supermix for RT-PCR (Bio-Rad Laboratories, Hercules, CA, USA), which utilizes a mix of random primers and oligo(dT), enabling reverse transcription of total RNA, including both polyadenylated and non-polyadenylated transcripts. RT-PCR was performed using synthesized cDNA. Each 10 µL reaction contained GoTaq® Green Master Mix (Promega), gene-specific primers (Table S1), nuclease-free water, and cDNA template. For *PIF3* amplification (∼180 bp), thermocycling conditions included an initial denaturation at 94 °C for 2 minutes, followed by 32 cycles of denaturation at 94 °C for 30 seconds, annealing at 60 °C for 45 seconds, and extension at 72 °C for 30 seconds, with a final extension at 72 °C for 5 minutes. The mitochondrial *CO1* gene (∼700 bp) was amplified under the following conditions: 94 °C for 2 minutes, followed by 30 cycles of 94 °C for 30 seconds, 55 °C for 30 seconds, and 72 °C for 45 seconds, with a final extension at 72 °C for 5 minutes. PCR products were resolved on 1% agarose gels stained with SYBR™ Green DNA gel stain and visualized under UV light. A 100 bp DNA ladder (for *PIF3*) and a 1 kb Plus DNA ladder (for *CO1*) were used as molecular size markers.

### Verification of viral protein production in passage 2 cells by Western blot

We assayed for the production of viral proteins to complement the DNA-and RNA-based detection of virus described above. We pooled equal volumes of each 240 hour cell control and virus infected passage 1 fluids from each cell line and separated a fraction of the pool by polyacrylamide gel electrophoresis (10%) under reducing conditions as described by Laemmli (1970). Then we transferred to an Immobilon membrane (Millipore) by western blotting using a Bio Rad Protean mini-cell with a lane of Chameleon 700 Pre-stained Protein Ladder (LI-COR, NE, USA). After transfer the membrane was blocked with Dulbecco’s Phosphate – Buffered Saline (DPBS, Corning 21-031-CV), 2% BSA and 0.1% Tween-20 for 1 hour at room temperature. The blocking solution was removed, and the membrane was soaked in anti-vp39 antibody (Schedl et al., 2025) diluted 1:2000 and mouse anti-Actin (JLA20-s, Developmental Studies Hybridoma Bank, IA, USA) diluted 1:500 in DPBS, 2% BSA, 10% fetal bovine serum and 0.1% Tween-20 and incubated for 1 hour at room temperature. Then, the membrane was washed 3x with DPBS followed by a 1-hour incubation in DPBS, 2% BSA and 0.1% Tween-20 containing Alexa Fluor 488 goat anti-rabbit (Invitrogen, A11008) diluted 1: 4000 and goat anti-mouse IRDye 680RD (LI-COR, NE, USA) diluted 1:5000. After incubation with the conjugates the membrane was washed 3x with DPBS and visualized with a Licor Odyssey M imaging system. The image was processed with FIJI using the Quick Figures plugin (Schindelin et al., 2012).

### Fluorescent focus-forming unit (FFU) assay to measure viral titer

FH1 and CCA1 inoculated on *D. innubila, D. virilis* and *D. melanogaster* cells for 240 hours were titrated using a fluorescent focus-forming unit (FFU) assay (Schedl et al., 2025). For simplicity, we pooled fluids from each replicate and inoculated in *D. virilis* (Dv-1) cells since a) the virus replicates well in these cells, and b) *D. innubila* cells are difficult to count. We also performed the FFU assay on the inoculum to calculate the increase in FFU over the course of the experiment.

Briefly, using 48-well plates, the interior 24 wells were seeded with *D. virilis* Dv-1 cells at approximately 10^5^/well in a 500µl Schneider’s *Drosophila* medium and incubated at 23°C for 24 hours for cell attachment. The surrounding wells were filled with DPBS to control for evaporation. After 24 hours, the culture medium was removed and 200µl viral fluid was inoculated onto cells. There were four replicate wells inoculated per dilution (10^-1^, 10^-2^,10^-3^, 10^-4^, and 10^-5^) and four wells that had no virus, the viral fluids were diluted in Schneider’s *Drosophila* medium. Plates were incubated for 48 hours at 23^0^C, then the culture media was removed and 200ul of chilled 80% acetone was added to each well, and incubated at-20^0^C for 30 minutes to fix cells. The acetone was then removed and wells were air dried for 20 minutes. 200ul of 2% BSA, 0.1% Tween in 1X DPBS blocking buffer solution was carefully added to each well and incubated at room temperature for 30 minutes. After incubation, the BSA solution was removed and 200μL of antibody solution (1:2000 dilution of P39 DiNV capsid protein primary anti-rabbit antibody in 2% BSA, 0.1% Tween in 1X DPBS) was added to each well, and the plates were incubated at 4^0^C overnight.

After incubation with the antibody, the plates were washed three times with DPBS then 200μL of secondary antibody solution (1:4000 dilution of goat anti-rabbit IgG Alexa Fluor 488 secondary antibody in 2% BSA DPBS) was added to each well and incubated for 1 hour in the dark. After incubation with the secondary antibody, the plates were washed three times with DPBS then imaged with a Leica Thunder microscope with the 10X objective in bright field and GFP channels. In total, there were six plates representing six different viral fluids (FH1 and CCA1 inoculated on Dinn-1, Dv-1 and S2 cells) tittered. FFU per milliliter was calculated by multiplying the average number of FFU by the dilution factor and dividing by the inoculation volume (mL). Increase in FFU was determined by dividing the FFU/ml after the experiment to that of the inoculum (adjusted for dilution in the experiment).

### Analysis

The raw Cq data was organized in Microsoft Excel and analyzed with R using the ggplot2 for figures, base R for most statistics, and DescTools for post-hoc statistical testing (Signorell, 2024; Team, 2021; Wickham, 2016). We used standard ANOVA, Tukey tests and Dunnett’s for most analyses. To be conservative, we decided to retain all outliers, assuming the sample replicates should swamp out the effect of rare outliers. Any Cq value that was undetermined after 32 cycles was converted to a 32 to reflect that the sample does not have sufficient target DNA in the sample to react with the primers after 32 cycles (Goni et al., 2009) and because Cq variation increases with decreasing target concentration (Forootan et al., 2017). We did not consider the primer efficiencies in the Cq calculations (Ruiz-Villalba et al., 2021) since the primer efficiencies are close to 100% for each primer set (Supp Table 7). Our experiment allowed for three different types of analyses all based on nucleic acids: virus per cell, virus per volume, and cells per volume, where volume represents the amplification of our extraction control Lambda DNA. In all cases, we perform statistics on the ΔΔCq values since they are technically log2 transformed. Plots either use the ΔΔCq value or the 2ΔΔCq value which can be interpreted as the relative amount of, say, viral DNA per host cellular DNA. For each passage 1 experiment and each comparison (cells per volume, virus per volume, virus per cell), we first performed a full ANOVA model to assess the effect of treatment, cell type, experimental block (if available) and the interaction of treatment and cell type on increase in the inoculated condition compared to the uninoculated control. For example: lm(Cq_cells_per_volume_0hour - Cq_cells_per_volume_240hour ∼ Cells x Treatment + Cells + Treatment + Block). Then we performed a Tukey test on that model examining the pairwise effects of cell type: TukeyHSD(aov(model), which=“Cells”), where model refers to the model described in the previous sentence. Finally, performed an ANOVA test using each cell type independently to test for a change compared to uninoculated controls in that specific cell type: lm(Cq_cells_per_volume_0hour - Cq_cells_per_volume_240hour ∼ + Infection + Block), but restricting to a single cell type. For the passage 2 experiment with passage 1 fluids from the fly homogenate treatment, our model included only treatment (which involved 5 inoculants and 1 mock infected control), followed by a Dunnett test to compare each treatment to the control, but otherwise is like what was described for the passage 1 experiment. For the cell-culture adapted, passage 2 experiments, we employed three different cell types, but all were considered independently using the model described above.

## Results

### Fly Homogenate Passage I – Naïve virus replicates its genome in D. innubila and D. virilis cells, but less so in D. melanogaster cells

When inoculated with homogenate of wild-caught, DiNV-infected flies, all three cell types grew over the ten days of infection with mock inoculated cells increasing their genomes 3.37-fold, 22.8-fold, and 36.7-fold in *D. innubila* (Dinn-1), *D. virilis* (Dv-1), and *D. melanogaster* (S2) cells, respectively (Fig 1A, Table S2). All factors (cell type, infection, cell type by infection interaction, and block) were significantly associated with changes in cells per volume (Table S3, p<0.01), and all three cell types were significantly different from each other for cell growth (Tukey test p<0.0001, Table S4). Taken independently, for Dinn-1 cells inoculated with virus, the host cell abundance increase was 4.64-fold, but this was not significantly different than the mock inoculated cells (p=0.063, Table S5). However, for Dv-1 and S2 cells, viral inoculation was associated with a negative effect on cell growth (Dv-1 p=0.002, S2 p=0.00002, Table S5). Thus, inoculation with DiNV appears to negatively impact growth in *D. virilis* (Dv-1) and *D. melanogaster* (S2 cells), but not in *D. innubila* (Dinn-1) cells.

**Figure 1.**
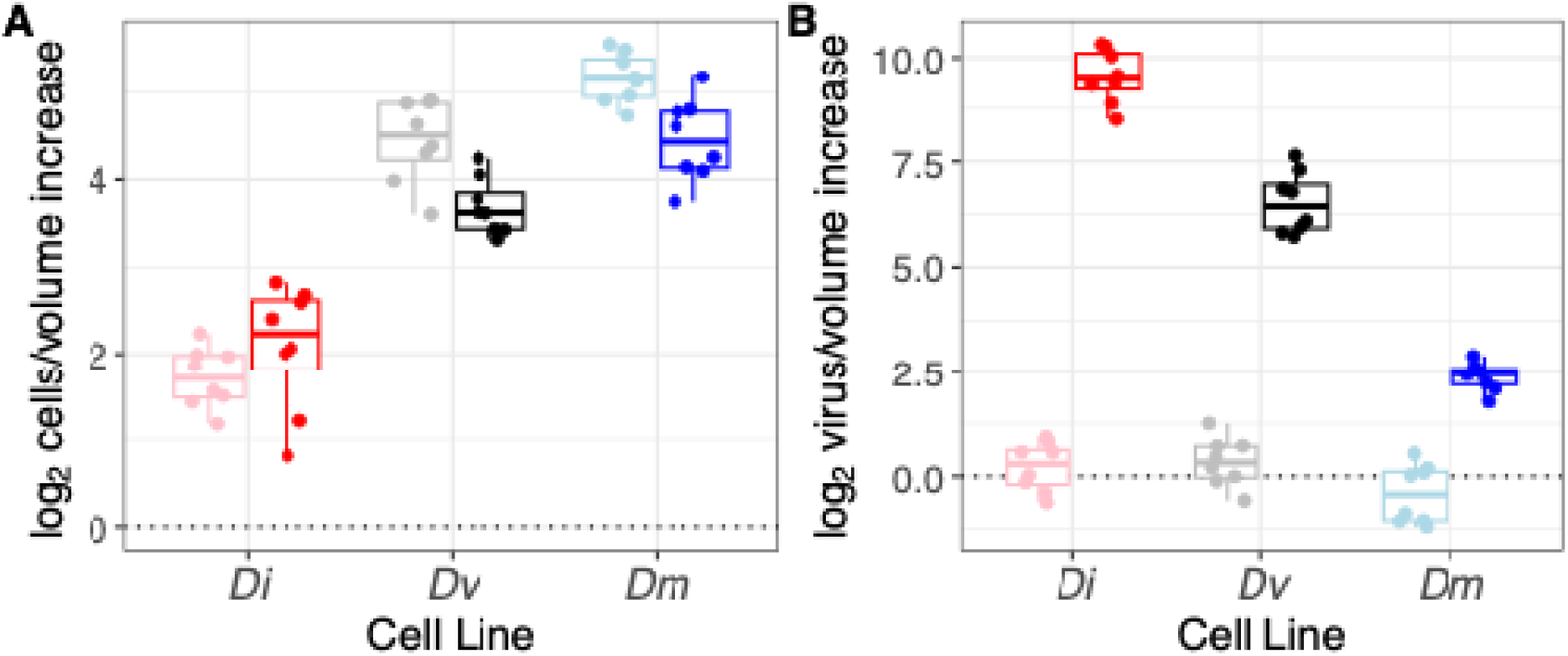
**Passage 1 with fly homogenate cell and virus log_2_ genome fold increase (**ΔΔ**Cq).** A) host cell genome increase for each of the three cell lines with and without virus inoculation. B) virus genome increase per volume for each of three cell types with and without viral inoculation. Cell types are: Dinn-1 (Di, *D. innubila*), Dv-1 (Dv, *D. virilis*) and S2 (Dm, *D. melanogaster*). Lighter colors are mock infected and darker colors are infected with virus.

**Figure 1.**
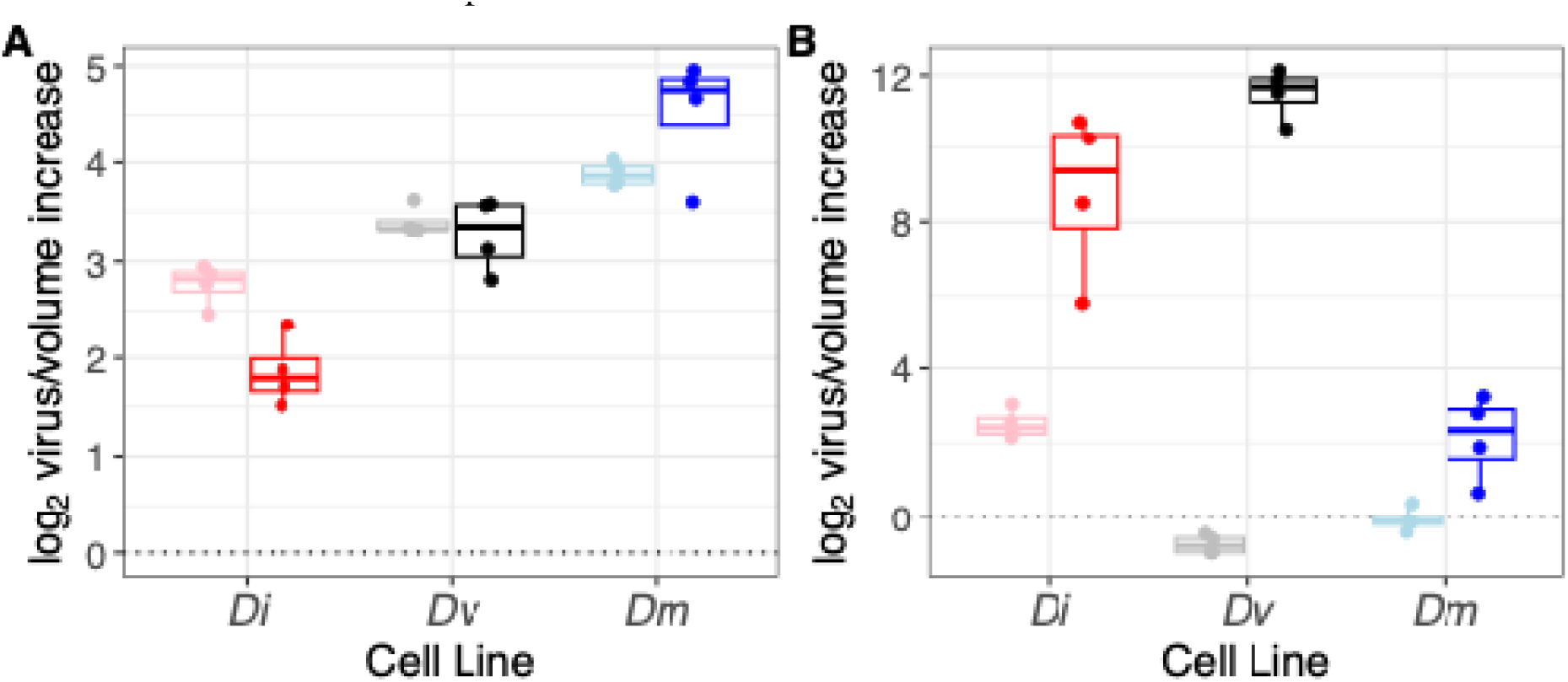
**Adapted virus Passage 1 cell and virus log_2_ genome fold increase (**ΔΔ**Cq**). a.) host cell genome increase for each of the three cell lines with and without virus inoculation. b.) virus genome increase per volume for each of three cell types with and without viral inoculation. Cell types are: Dinn-1 (Di, *D. innubila*), Dv-1 (Dv, *D. virilis*) and S2 (Dm, *D. melanogaster*). Lighter colors represent mock inoculated and darker colors represent DiNV inoculated.

**Figure 2.**
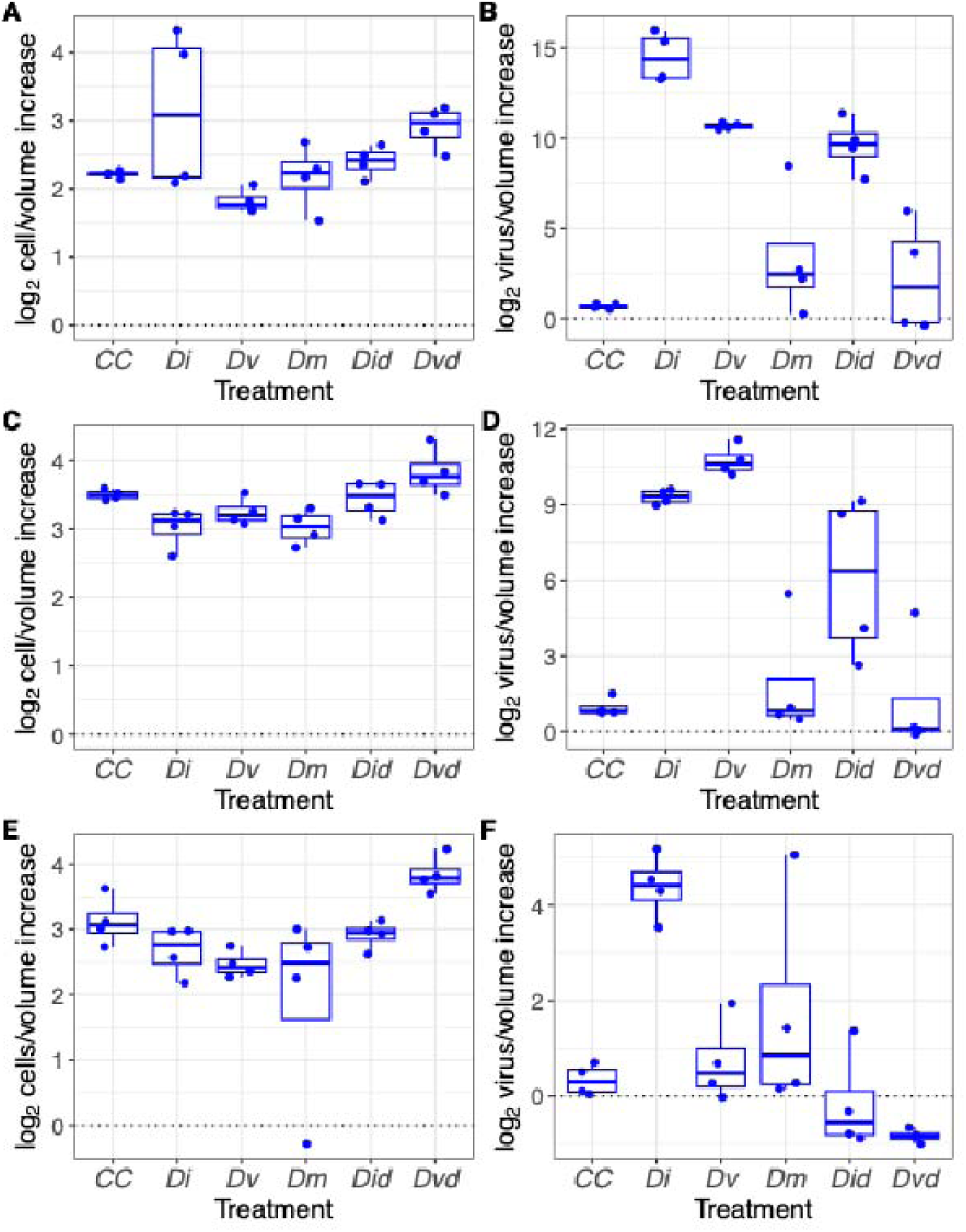
**Adapted Passage 2 cell and virus log_2_ genome fold increase (**ΔΔ**Cq).** *A*) host Dinn-1 cell genome increase per volume with and without virus inoculation. *B*) virus genome increase per volume with and without viral inoculation *in Dinn-1 cells*. *C*) host D*v*-1 cell genome increase per volume with and without virus inoculation. *D*) virus genome increase per volume with and without viral inoculation *in Dv-1 cells*. *E*) host *S2* cell genome increase per volume with and without virus inoculation. *F*) virus genome increase per volume with and without viral inoculation *in S2 cells*. Treatment is passage 1 virus derived from the passage 1 study (mock = CC, Di from *D. innubila*, Dv from *D. virili*s, Dm from *D. melanogaster*, Did from diluted *D. innubila*, Dvd from diluted *D. virilis*.

Ignoring host nucleic acids, the DiNV genome copy number per volume (where Lambda DNA extraction controls are used as a proxy for volume) increased 809-fold, 103-fold and 5.35-fold in Dinn-1, Dv-1, and S2 cells respectively (Figure 1B, Table S2). The full statistical model shows a significant effect of all factors (cell type, treatment, block, and cell type by treatment interaction) on DiNV genome copy per volume (p<0.00001, Table S6), and all cell types were significantly different in terms of viral increase per volume (Tukey test p<0.00001, Table S7). In all three cases, however, the viral genome increase was significant (Dinn-1 p<0.00001, Dv-1 p<0.00001, S2 p<0.00001, Table S8).

DiNV genome copy number per cell was qualitatively similar to the results for virus per volume. DiNV genome copy number per cell increased 190-fold, 8-fold and 0.24-fold in Dinn-1, Dv-1 and S2 cells respectively. None of the mock infected cells showed a virus genome increase. Note that the decrease in virus in S2 makes sense: S2 cells grow well in culture and DiNV does *not* grow well in S2 cells, so the viral DNA copy number relative to the host cell DNA copy number actually decreased. The full statistical model again shows a significant effect of all factors (cell type, treatment, block, and cell type by treatment interaction) on DiNV genome copy per cell (p<0.00001, except block is more borderline p=0.0252, Table S9, Figure S1). All cell types were again significantly different in terms of viral increase per volume (Tukey test p<0.00001, Table S10) and taken individually, each cell line showed a significant increase in viral genome copies per cell compared to the controls (p<0.00001, Table S11).

In this first virus growth experiment we found that DiNV derived from wild caught fly homogenate was able to replicate its genome in the *Drosophila* cells for at least one virus passage. The cells derived from *Drosophila innubila* are the most permissive followed by *Drosophila virilis* and the least permissive are the S2 cells that are derived from *Drosophila melanogaster*. Thus, for our limited sample, cells derived from more closely related hosts (*D. innubila* Dinn-1 and *D. virilis* Dv-1) are permissive to viral genome replication than those more distantly related (*D. melanogaster,* S2). However, viral genome replication does not mean those infections produced infectious virions.

### Fly homogenate Passage 2 – Minimal viral genome replication from naive passage 1 virus

To determine if infectious DiNV was generated in the Fly Homogenate Passage 1 growth study in any of the cells, we inoculated a new set of Dinn-1 cells with the 240-hour, virus positive Passage 1 fluids and we included 2 additional groups that consisted of fluids collected from the Dinn-1 and Dv-1 groups that were diluted to have approximately an equal DiNV genome copy number compared to the S2 group, in an attempt to equilibrate inoculum doses. Surprisingly, none of the virus fluids from passage 1 showed much evidence of growth when reinoculated. In fact, while most of the cells grew well (Figure 2A), none of the inoculants showed genome replication significantly different from the mock inoculated cell control group (Figure 2B).

**Figure 2.**
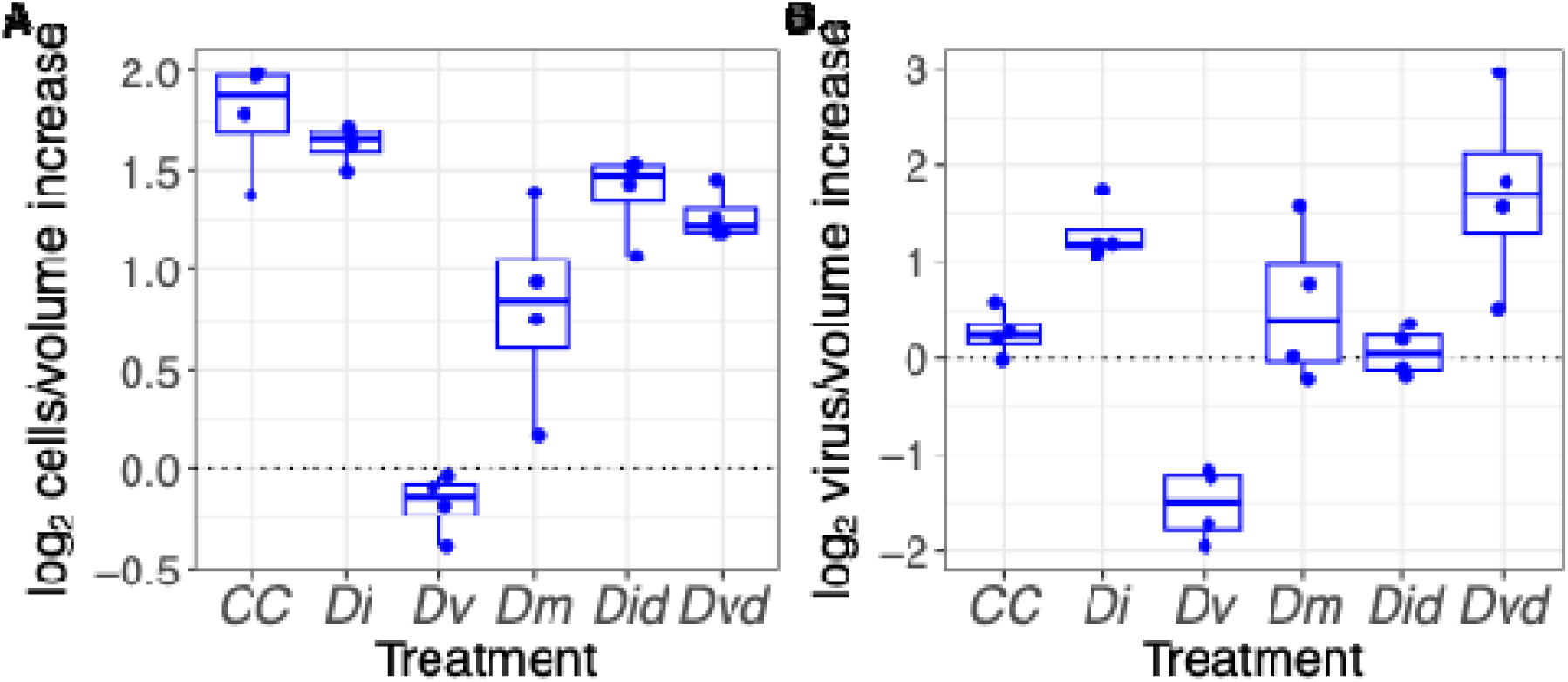
Passage 2 from fly homogenate cell and virus log_2_ genome fold increase (**ΔΔ**Cq). A) host Dinn-1 cell genome increase per volume with and without virus inoculation. b.) virus genome increase per volume with and without viral inoculation. Cell type is Dinn-1. Treatment is passage 1 virus derived from the passage 1 study (mock = CC, Di from *D. innubila*, Dv from *D. virilis*, Dm from *D. melanogaster*, Did from diluted *D. innubila*, Dvd from diluted *D. virilis*.

In mock inoculated Dinn-1 cells the cell genome per volume mean fold increase was 3.48-fold (Table S12). While there was an overall effect of treatment on cell growth (p<0.00001, Table S13), the only treatments that showed significant differences from the control were cells inoculated with Dv-1 fluids (0.89-fold increase; Dunnett Test p<0.00001, Table S14, Fig 2A), and S2 cells (1.83-fold increase; Dunnett’s Test, p=0.00026, Table S14, Fig 2A).

All samples showed very little viral genome replication. The virus per volume genome increase of the Dinn-1 mock inoculated cell control was 1.2-fold (Table S12). Overall, there was a significant effect of inoculant on viral genome increase per volume (p<0.00001, Table S15, Fig 2B). Of the three undiluted inoculants, only Dv-1 derived virus and the diluted Dv-1 virus was significantly different than the cell control and it showed a difference in viral copies (Dunnett’s test, Dv-1 p=0.0017, Dv-1 diluted p=0.0093, Table S16, Fig 2B). Interestingly, Dv-1-derived virus shows *less* viral genome increase compared to controls.

The pattern was similar for virus per cell. There was an overall significant effect of treatment on change in viral genomes per cell (p=0.00535, Table S17, Figure S2). The Dinn-1 mock inoculated cell control had a virus genome per cell increase of 0.349-fold (Table S12). The passage 1 virus inoculum derived from the Dinn-1, Dv-1, S2, and diluted Dinn-1 did not produce a significant virus genome per cell increase compared to the mock control, but the diluted Dv-1 passage 1 fluid produced a significant virus genome increase compared to the mock infected Dinn-1 cell control (1.66-fold, Dunnett’s Test p= 0.0046, Figure S2, Table S18).

We conclude that the virus isolated from wild caught flies and passaged once through two cell types (Dinn-1 and S2) was unable to form enough infectious virions to be passaged to new cells, even if the cells are from the focal host.

### Cell culture adapted virus Passage 1 – adapted viral genome replicates better than naïve virus

We isolated virus after incubating with *D. innubila* cells for about one year (Schedl et al., 2025). We refer to this virus as cell-culture adapted virus. Preliminary experiments with the adapted virus demonstrated that the virus isolate was able to produce progeny virions in the Dinn-1 cells, so we repeated both the of the previous growth studies with this adapted virus.

The cells in the adapted virus trials grew similarly to the fly homogenate passage 1 trials. The mock inoculated host cell genome copy per volume mean fold increase was 6.84-fold, 10.56-fold and 14.76-fold for Dinn-1, Dv-1 and S2 cells, respectively (Table S19). Overall, both cell type (p<0.00001) and cell type by treatment interaction (p=0.00142) were significantly associated with change in cell genome copy per volume, but treatment alone was not (p=0.37, Table S20, Fig 3A). Furthermore, all three cell types grew significantly differently from each other (Tukey test p<0.0001, Table S21). The growth of the inoculated Dinn-1 cells was significantly lower when considered alone, compared to the mock infected Dinn-1 cells (p=0.0048, Table S22, Figure 3A), but inoculation was not associated with differences in cell growth for Dv-1 (p=0.55) or S2 cells (p=0.0925).

For viral genome copy per volume, cell type, treatment and their interaction were all significantly associated with change over time (p<0.00001, Table S23, Figure 3b). The virus genome copy number per volume mean increase was 839.41-fold, 3145.47-fold and 5.38-fold in Dinn-1, Dv-1 and S2 cells, respectively. Dinn-1 and Dv-1 cells both showed significant pairwise differences from S2 (p<0.00001, Table S24), but not were not different from each other (p=0.84126). These increases were all significant compared to mock infected cells, a measure of background levels of amplification (Dinn-1 p=0.00057, Dv-1 p<0.00001, S2 p=0.00976, Table S25, Fig 3B).

Finally, when analyzing viral genome increase per cell, cell type, treatment and cell type by treatment interaction were all significant (p<0.00001, Table S26, Fig S3), and Dinn-1 and Dv-1 were significantly different than S2 (p<0.00001, Table S27) but not from each other (p=0.08321). In virus inoculated Dinn-1 and Dv-1 cells the virus genome per cell mean fold increase was 233-fold and 319-fold, respectively (Table S19), but only 0.07-fold in S2 cells. The virus increase in the inoculated Dinn-1 and Dv-1 cells was significant compared to the mock inoculated cells (Dinn-1 p=0.00057, Dv-1 p<0.00001, Table S28, Fig S3), but not in S2 cells (p=0.110).

### Cell Culture Adapted Passage II – Adapted virus produces infectious virions and can be passaged

To test whether the adapted virus produced infectious virions in cell culture, we inoculated Dinn-1, Dv-1 and S2 cells with pooled virus from the cell culture adapted Passage 1 growth study. The virus inoculum was prepared from the 240-hour samples from each virus treatment group in the passage 1 study and we included a diluted Dinn-1 Passage 1 and diluted Dv-1 Passage 1 group (the virus fluid in these groups was diluted to have equivalent Cq values as the S2 group). Unlike the fly homogenate passage 2 study, we included all three cell types (Dinn-1, Dv-1 and S2) with 5 different virus preparations (Dinn-1/DiNV P1, Dv-1/DiNV P1, S2/DiNV P1, diluted Dinn-1/DiNV P1 and diluted Dv-1/DiNV P1). For this experiment, we elected to analyze infections in each cell type independently.

The host cell (*D. innubila*) genome per volume mean fold increase was 4.64-fold, 11.5-fold, 3.55-fold, 4.69-fold, 5.33-fold, and 7.61-fold for fluids from mock infected, Dinn-1, Dv-1, S2, diluted Dinn-1 and diluted Dv-1, respectively (Table S29). There was a significant effect of treatment on cell growth per volume (p=0.0275, Table S30, Fig 4A), but none of the inoculated cells grew significantly differently than the mock infected cells (Dunnett’s test p>0.05, Table S31, Fig 4A).

The virus genome copy number per volume mean fold increase in Dinn-1 cells inoculated with passage 1 fluids was 31,1567-fold, 1631-fold, 90-fold, 1122-fold, and 19-fold for fluids from Passage 1 Dinn-1, Dv-1, S2, diluted Dinn-1 and diluted Dv-1, respectively (Table S29, Figure 4B). Treatment was significant in the overall model (p<0.00001, Table S30, Fig 4B), and fluids from passage 1 Dinn-1 cells (p<0.00001, Table S31), diluted Dinn-1 cells (p=0.00004) and Dv-1 cells (p<0.00001) were significantly different from controls, but neither fluids from passage 1 S2 cells (p=0.2673) nor diluted Dv-1 cells (p=0.529) were different from mock infected controls. So, while diluted Dinn-1 fluids from passage 1 led to significant viral growth, diluted Dv-1 fluids did not, even though both undiluted fluids did show significant growth.

The virus genome per cell increase in Dinn-1 cells inoculated with mock inoculation or passage 1 fluids from the Dinn-1, Dv-1, S2, diluted Dinn-1 and diluted Dv-1 groups was 0.34-fold, 2622-fold, 465-fold, 14-fold, 201-fold and 3-fold, respectively (Table S29, Fig S4). Treatment had a significant effect on virus genome increase per cell (p<0.00001, Table S30), and the Dinn-1, Dv-1 and diluted Dinn-1 inoculation groups had significant virus growth compared to controls (Dunnett’s Test p<0.00001, Table S31, Fig S4), while fluids from S2 and diluted Dv-1 did not (p=0.2296 and p=0.9529, respectively, Table S31).

Results for Dv-1 cells were remarkably like those for Dinn-1 cells, with a significant effect of treatment for cell growth but no differences in any samples from controls (Table S32, Table S33, Fig 4C).

Similarly, for both virus per cell and virus per volume increase fluids from Dinn-1, Dinn-1 diluted, and Dv-1 led to significant increases, but not S2 and Dv-1 diluted did not (Table S33, Fig 4D, Fig S5). In S2 cells, like the others there was a significant effect of treatment on cell growth but no differences in any samples compared to the controls (Table S34, Table S35, Fig 4E). The only inoculant that showed any growth in S2 cells was from Dinn-1 passage 1 fluids (p=0.0004 for virus per volume, p=0.0048 for virus per cell, Table S34, Table S35, Fig 4F, Fig S4).

### Further evidence that DiNV has adapted to cell culture

We reasoned that, given the adapted virus but not the naïve virus produced infectious virus in *D. innubila* and *D. virilis* cells, we should only see evidence of viral gene transcription in those cells. To that end, we extracted RNA from each of our passage 2 samples and performed RT-PCR using primers for the viral *PIF3* transcript and for host *COI*. As expected, only the 240-hour *D. innubila* and *D. virilis* samples from the adapted virus showed evidence of viral transcription, but all samples showed evidence of *COI* transcription (Fig S5).

Finally, we aimed to confirm the production of viral proteins in passage 2 samples by Western blot. Surprisingly, only the *D. virilis* cells inoculated with cell adapted virus were positive when assayed with anti-vp39 antibody (Fig S6). None of the controls nor the cells inoculated with naïve virus were positive. This is unexpected because *D. innubila* and *D. virilis* cells showed similar increases in genome copy number for cell adapted virus and both were also positive for viral transcripts. However, it is possible that the amount of vp39 protein from *D. innubila* was just below detection thresholds.

Overall, the cell-adapted virus grew well in *D. innubila* and *D. virilis* cells and less well in *D. melanogaster* cells, suggesting that at least two cell types have promise for future experiments.

### FFU assay confirms differences between adapted and ancestral virus

To confirm the qPCR results, we performed FFU assays on inoculum and viral fluids after the first passage. The naïve fly homogenate virus grown in all three cell lines showed very little increase (8.56x, 6.52x and 4.91x for *D. innubila*, *D. virilis*, and *D. melanogaster* cells respectively, Fig 5). Note, that while slight, all of these increases are significantly greater than 0 (T-test p-value <0.05). The adapted virus, however increased substantially in both *D. innubila* (656x) and *D. virilis* (646x), but not *D. melanogaster* (4.08x). In fact, the increase for *D. melanogaster* after inoculation with adapted virus was not significantly different from the increase after inoculation with naïve virus (T-test p-value=0.15). In a full ANOVA model, there was an effect of cell type, virus (adapted or naïve) and the interaction between cell type and virus (p<0.0001 for all factors).

**Figure 5.**
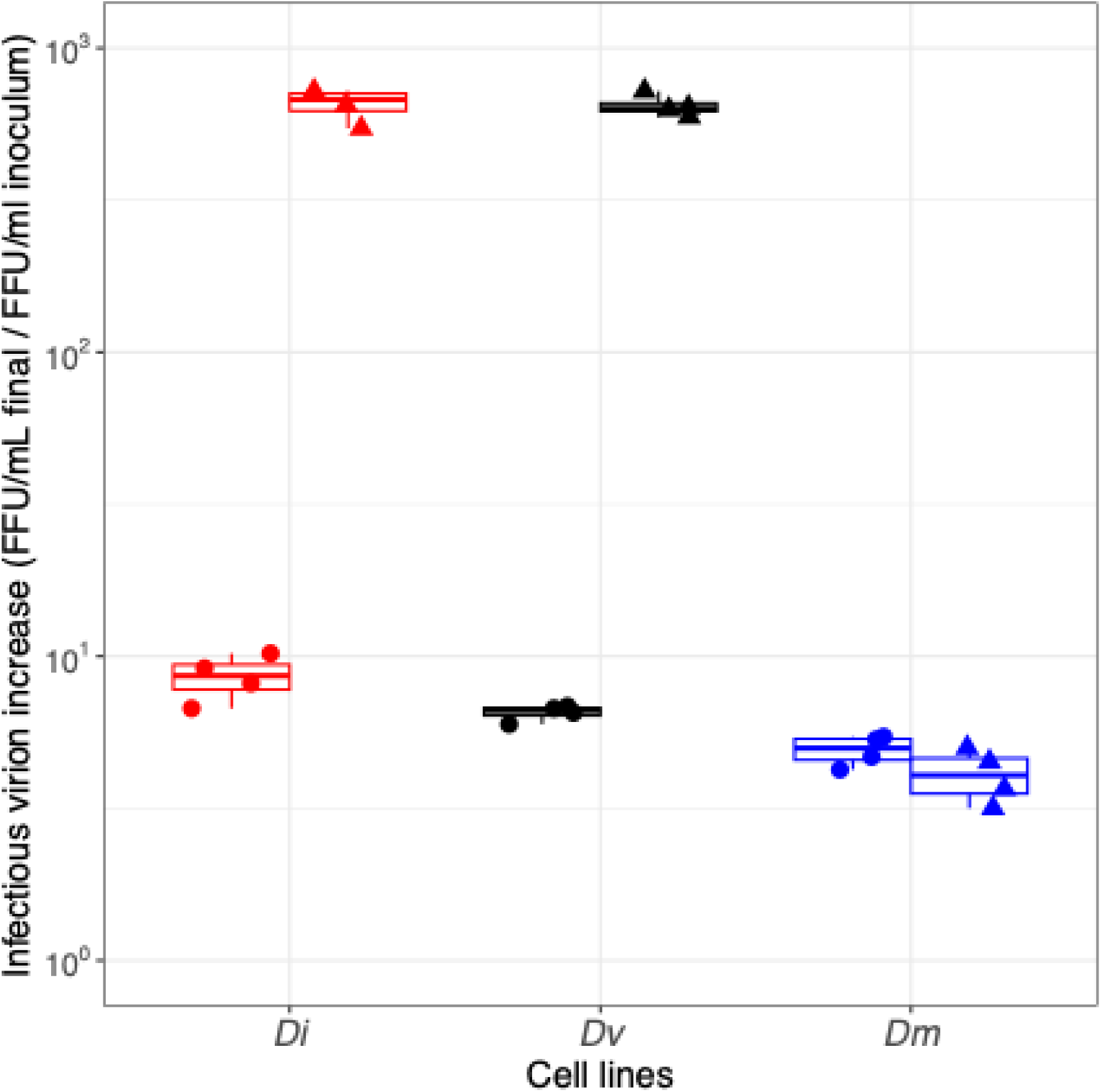
Infectious virions increased during the first passage but much more for the adapted virus than the naïve. A fluorescent focus-forming unit assay was performed using *D. virilis* (Dv-1) cells. Values represent the relative increase in viral titer (FFU/ml) from inoculation to the end of the experiment. (Di from *D. innubila*, Dv from *D. virili*s, Dm from *D. melanogaster*, circles represent naïve virus and triangles represent adapted virus).

## Discussion

Viruses are among the most abundant biological entities on earth (Iranzo et al., 2016; Koonin and Dolja, 2013; Krupovic et al., 2019; Lezcano et al., 2023) and the main activity of an individual virus is to infect a cell and replicate, then the progeny will move to a new host to repeat the cycle (David M. Knipe, 2013). Some viruses are generalists like the baculovirus *Autographa californica* multiple nucleopolyhedrovirus (AcMNPV) which will infect cells from at least 30 species of *Lepidoptera* (Iranzo et al., 2016; Krupovic et al., 2019). Others are very specific about the type of host cell they can infect and reproduce in, like variola virus which only infects susceptible humans (Lezcano et al., 2023). Viral host range is an important characteristic to understand when considering the virus life cycle and potential spillover to new hosts. The development of cell-culture infection models depends on an understanding of this host range.

To gain an understanding of the infection range of DiNV we studied the infection specificity of *Drosophila innubila* Nudivirus in *Drosophila* cell culture and the impact, if any, of virus infection on the growth of the inoculated cells. We used both a naïve virus isolate, and a virus isolate that had adapted to grow in cell culture. For each virus, we performed two growth experiments: an initial inoculation (passage 1) and an attempt to passage the virus after initial growth (passage 2) in cells from 3 species of *Drosophila*. We found that naïve DiNV derived from wild caught fly homogenate had variable ability to replicate in the three cell lines, but very little ability to be passaged. In contrast, the cell-culture adapted virus grew well initially (passage 1) but was also able to reinfect (passage 2) new cells. Overall, we found that cells derived from species more closely related to the focal host, *D. innubila* were better hosts for the virus, but that in some ways, the *D. virilis* (Dv-1) cells were comparable to the *D. innubila* (Dinn-1) cells.

The effects of viral inoculation were inconsistent across cell type and across the two passage one experiments (using naïve virus and cell culture adapted virus). With naïve virus, both *D. virilis* Dv-1 and *D. melanogaster* S2 cells seemed to grow more poorly than *D. innubila* Dinn-1. However, when inoculated with the adapted virus, Dinn-1 cells grew slightly less than mock and S2 cells grew slightly better. The inconsistency of the results between experiments and the relatively modest effect of inoculation makes us hesitant to draw strong conclusions, but it does appear that under some conditions minor cytopathic effect may occur in *some* cell lines.

Surprisingly, although the naïve virus from fly homogenate replicated its genome well in both *D. innubila* Dinn-1 and *D. virilis* Dv-1 cells, the fluids harvested from those cells did not contain viruses that were able to reinfect cells in a second passage. This contrasts with the cell culture-adapted virus which grew well in both passage 1 and passage 2 upon reinoculation. Future work will aim to discover the genetic changes in the virus that enabled it to be passaged through cell culture. We speculate that either cellular components or environments were missing necessary components for viral production or that the 10-day experimental course was too short to produce infectious virions from the naïve virus. Other nudiviruses and baculoviruses utilize host cell components (proteins) for viral assembly and even as parts of the infectious virions (Bezier et al., 2017; Hou et al., 2013). If DiNV also is dependent on host proteins or other structures for complete viral assembly, the three cell types employed may be missing those components. This is true even of the native host Dinn-1 cells if those cells mimic host tissues that are not conducive to viral assembly. Alternatively, our experiment may have ended too soon for full viral particle assembly which would render whatever viral nucleic acids and proteins had assembled non-infectious.

Both scenarios are consistent with ample nucleic acid replication and production of protein without producing infectious virions.

## Conclusion

We inoculated cells from 3 species of *Drosophila* with 2 strains of *Drosophila innubila* Nudivirus. We found that cells derived from *Drosophila innubila* replicate both virus strains to high genome copies and they appear to be the most permissive of the three to virus infection by either strain of DiNV. The two strains of virus appear to have some differences as the virus derived from wild caught flies will infect the three cells but does not appear to produce infectious virus progeny. The cell culture-adapted virus strain is infectious when inoculated on Dinn-1 and Dv-1 cells but not on S2 cells. This cell culture-adapted virus will produce infectious progeny which allows for the ability to grow a sufficient volume of virus of which can be used for host/pathogen genetic conflict studies in the context of a natural DNA virus infection in *Drosophila*.

## Acknowledgements

We thank Margaret E. Schedl, Brian D. Ackley, Robin C. Orozco and members of the Unckless lab for valuable feedback. We thank Dr. Kausik Si and the Drosophila Genomics Resource Center (NIH grant 2P40OD010949) for cell lines. Funding was provided by NSF EAGER 2135167 to RLU.

