## Supplemental Material for "Specificity of *Drosophila innubila* Nudivirus Infection in *Drosophila* Cell Culture"

### Supplemental Materials

### Supplemental Tables

| **Gene** | **Use** | **Forward** | **Reverse** | **R^2^** | **Slope** | **Eff.** |
| --- | --- | --- | --- | --- | --- | --- |
| PIF3 | DiNV | AGCTGGCGACACGAATAAAATA | CCGGGAGCATTTAAAATATGAG | 0.999 | -3.33 | 0.998 |
| Lambda | Spike | CGGCGTCAAAAAGAACTTCC | GCATCCTGAATGCAGCCATA | 0.994 | -3.45 | 0.948 |
| Tpi | Dinn-1 | CCCAATCGGCGCCAAT | GCCGGACAGAATGCCTACAA | 0.991 | -3.22 | 1.044 |
| RPL11 | Dv-1 | GCAGCCCGTGTTTTCTAAGG | TACTCGCGAACTTTCAAGCCAC | 0.993 | -3.53 | 0.922 |
| Rp49 | S2 | AGCATACAGGCCCAAGATCG | TGTTGTCGATACCCTTGGGC | 0.998 | -3.18 | 1.063 |
| COI | Host RT-PCR | GGTCAACAAATCATAAAGATATTGG | TAAACTTCAGGGTGACCAAAAAATCA | NA | NA | NA |

**Table 2.S1. Primers used in this study and their qPCR efficiencies.** Eff = efficiency. Note that the PIF3 primers were used for both qPCR and RT-PCR.

|  | **Cells/ Volume** | **Log_2_ Cells/ Volume** | **Virus/**  **Cell** | **Log_2_ Virus/ Cell** | **Virus/ Volume** | **Log_2_ Virus/ Volume** |
| --- | --- | --- | --- | --- | --- | --- |
| Dinn-1 Control | 3.368 | 1.174 | 0.36 | -1.493 | 1.247 | 0.221 |
| Dinn-1 DiNV | 4.637 | 2.075 | 190.49 | 7.461 | 809.16 | 9.536 |
| Dv-1 Control | 22.804 | 4.447 | 0.063 | -4.113 | 1.350 | 0.334 |
| Dv-1 DiNV | 13.174 | 3.687 | 7.816 | 2.834 | 102.836 | 6.52 |
| S2 Control | 36.706 | 5.171 | 0.022 | -5.585 | 0.829 | -0.414 |
| S2 DiNV | 22.926 | 4.452 | 0.244 | -2.067 | 5.352 | 2.385 |

**Table 2.S2. Fly homogenate passage 1 experiment**. Results are reported as either ΔΔCq (e.g. Log_2_ Cells/ Volume) or 2^ΔΔCq^ (e.g. Cells/Volume). Inoculum source is homogenized DiNV-infected wild-caught *D. innubila.*

|  | **Df** | **Sum Sq** | **Mean Sq** | **F-value** | **P-value** |
| --- | --- | --- | --- | --- | --- |
| Cells | 2 | 73.50 | 36.75 | 214.401 | <0.00001 |
| Treatment | 1 | 1.67 | 1.67 | 9.746 | 0.003289 |
| Block | 1 | 1.96 | 1.96 | 11.459 | 0.001578 |
| Cells:Treatment | 2 | 3.23 | 1.62 | 9.432 | 0.000427 |
| Residuals | 41 | 7.03 | 0.17 |  |  |

**Table 2.S3. Fly homogenate passage 1 experiment for cell growth, full model ANOVA results.**

|  | **diff** | **lwr** | **upr** | **Adjusted P** |
| --- | --- | --- | --- | --- |
| Dm-Di | 2.916875 | 2.560930 | 3.2728203 | <0.00001 |
| Dv-Di | 2.172500 | 1.816555 | 2.5284453 | <0.00001 |
| Dv-Dm | -0.744375 | -1.100320 | -0.3884297 | 2.5e-05 |

**Table 2.S4. Tukey test results for cell type from fly homogenate passage 1 experiment for cell growth.**

| **Cell Type** | **Factor** | **Df** | **Sum Sq** | **Mean Sq** | **F-value** | **P-value** |
| --- | --- | --- | --- | --- | --- | --- |
| Dinn-1 | Treatment | 1 | 0.5202 | 0.5202 | 4.128 | 0.063141 |
|  | Phase | 1 | 2.8773 | 2.8773 | 22.832 | 0.000361 |
|  | Residuals | 13 | 1.6383 | 0.1260 |  |  |
| Dv-1 | Treatment | 1 | 2.3142 | 2.3142 | 14.597 | 0.00212 |
|  | Phase | 1 | 0.2475 | 0.2475 | 1.561 | 0.23352 |
|  | Residuals | 13 | 2.0610 | 0.1585 |  |  |
| S2 | Treatment | 1 | 2.0700 | 2.0700 | 40.87 | 2.37e-05 |
|  | Phase | 1 | 1.5098 | 1.5098 | 29.81 | 0.00011 |
|  | Residuals | 13 | 0.6585 | 0.0507 |  |  |

**Table 2.S5. Individual ANOVA testing for a treatment effect of viral inoculation on cell growth with each cell type treated separately.**

|  | **Df** | **Sum Sq** | **Mean Sq** | **F-value** | **P-value** |
| --- | --- | --- | --- | --- | --- |
| Cells | 2 | 123.9 | 61.9 | 284.16 | <0.00001 |
| Treatment | 1 | 446.6 | 446.6 | 2048.57 | <0.00001 |
| Phase | 1 | 6.6 | 6.6 | 30.13 | <0.00001 |
| Cells:Treatment | 2 | 85.0 | 42.5 | 194.85 | <0.00001 |
| Residuals | 418.9 | 0.2 |  |  |  |

**Table 2.S6. Fly homogenate passage 1 experiment for viral increase per volume, full model ANOVA results.**

|  | **diff** | **lwr** | **upr** | **Adjusted P** |
| --- | --- | --- | --- | --- |
| Dm-Di | -3.893437 | -4.294841 | -3.492034 | <0.00001 |
| Dv-Di | -1.450938 | -1.852341 | -1.049534 | <0.00001 |
| Dv-Dm | 2.442500 | 2.041097 | 2.843903 | <0.00001 |

**Table 2.S7. Tukey test results for cell type from fly homogenate passage 1 experiment for viral increase per volume**.

| **Cell Type** | **Factor** | **Df** | **Sum Sq** | **Mean Sq** | **F-value** | **P-value** |
| --- | --- | --- | --- | --- | --- | --- |
| Dinn-1 | Treatment | 1 | 347.1 | 347.1 | 1082.369 | <0.00001 |
|  | Phase | 1 | 1.2 | 1.2 | 3.646 | 0.0785 |
|  | Residuals | 134.2 | 0.3 |  |  |  |
| Dv-1 | Treatment | 1 | 153.11 | 153.11 | 721.62 | <0.00001 |
|  | Phase | 1 | 3.22 | 3.22 | 15.16 | 0.00185 |
|  | Residuals | 13 | 2.76 | 0.21 |  |  |
| S2 | Treatment | 1 | 2.0700 | 2.0700 | 40.87 | <0.00001 |
|  | Phase | 1 | 1.5098 | 1.5098 | 29.81 | 0.00011 |
|  | Residuals | 13 | 0.6585 | 0.0507 |  |  |

**Table 2.S8. Individual ANOVA testing for a treatment effect of viral inoculation on viral increase with each cell type treated separately.**

|  | **Df** | **Sum Sq** | **Mean Sq** | **F-value** | **P-value** |
| --- | --- | --- | --- | --- | --- |
| Cells | 2 | 371.6 | 185.8 | 743.176 | <0.00001 |
| Treatment | 1 | 502.9 | 502.9 | 2011.724 | <0.00001 |
| Phase | 1 | 1.3 | 1.3 | 5.394 | 0.0252 |
| Cells:Treatment | 2 | 60.4 | 30.2 | 120.892 | <0.00001 |
| Residuals | 41 | 10.2 | 0.2 |  |  |

**Table 2.S9. Fly homogenate passage 1 experiment for viral increase per cell, full model ANOVA results.**

|  | **diff** | **lwr** | **upr** | **Adjusted P** |
| --- | --- | --- | --- | --- |
| Dm-Di | -6.810312 | -7.240150 | -6.380475 | <0.00001 |
| Dv-Di | -3.623437 | -4.053275 | -3.193600 | <0.00001 |
| Dv-Dm | 3.186875 | 2.757037 | 3.616713 | <0.00001 |

**Table 2.S10. Tukey test results for cell type from fly homogenate passage 1 experiment for viral increase per cell.**

| **Cell Type** | **Factor** | **Df** | **Sum Sq** | **Mean Sq** | **F-value** | **P-value** |
| --- | --- | --- | --- | --- | --- | --- |
| Dinn-1 | Treatment | 1 | 320.7 | 320.7 | 1564.548 | <0.00001 |
|  | Phase | 1 | 0.4 | 0.4 | 1.845 | 0.197 |
|  | Residuals | 132.7 | 0.2 |  |  |  |
| Dv-1 | Treatment | 1 | 193.07 | 193.07 | 2800.06 | <0.00001 |
|  | Phase | 1 | 5.25 | 5.25 | 76.14 | <0.00001 |
|  | Residuals | 130.90 | 0.07 |  |  |  |
| S2 | Treatment | 1 | 49.53 | 49.53 | 280.448 | <0.00001 |
|  | Phase | 1 | 0.11 | 0.11 | 0.635 | 0.44 |
|  | Residuals | 132.30 | 0.18 |  |  |  |

**Table 2.S11. Individual ANOVA testing for a treatment effect of viral inoculation on viral increase per cell with each cell type treated separately.**

|  | **Cell/ Volumes** | **Log_2_ Cells/ Volume** | **Virus/**  **Cell** | **Log_2_ Virus/ Cell** | **Virus/ Volume** | **Log_2_ Virus/ Volume** |
| --- | --- | --- | --- | --- | --- | --- |
| Dinn-1 Control | 3.476 | 1.778 | 0.349 | -1.525 | 1.200 | 0.253 |
| Dinn-1 FH1 fluids | 3.980 | 1.629 | 0.808 | -0.333 | 2.500 | 1.299 |
| Dv-1 Control | 0.890 | -0.175 | 0.398 | -1.353 | 0.356 | -1.527 |
| Dv-1 FH1 fluids | 1.834 | 0.810 | 1.122 | -0.280 | 1.630 | 0.531 |
| S2 Control | 2.635 | 1.386 | 0.405 | -1.322 | 1.057 | 0.064 |
| S2 FH1 fluids | 2.419 | 1.270 | 1.66 | 0.445 | 3.925 | 1.715 |

**Table 2.S12. Fly homogenate passage 2 experiment.** Results are reported as either ΔΔCq (e.g. Log_2_ Cells/ Volume) or 2^ΔΔCq^ (e.g. Cells/Volume). Inoculum source is from fly homogenate passage 1 experiment from the Dinn-1 cells.

|  | **Df** | **Sum Sq** | **Mean Sq** | **F-value** | **P-value** |
| --- | --- | --- | --- | --- | --- |
| Treatment | 5 | 10.227 | 2.0454 | 28.5 | <0.00001 |
| Residuals | 18 | 1.292 | 0.0718 |  |  |

**Table 2.S13. ANOVA test results for Fly Homogenate passage 2 experiment, effect of treatment on cell growth.**

|  | **Diff** | **Lwr** | **Upr** | **Adjusted P** |
| --- | --- | --- | --- | --- |
| Di-CC | -0.14875 | -0.6719856 | 0.37448559 | 0.89332 |
| Did-CC | -0.39125 | -0.9144856 | 0.13198559 | 0.18539 |
| Dm-CC | -0.96625 | -1.4894856 | -0.44301441 | 0.00026 |
| Dv-CC | -1.95250 | -2.4757356 | -1.42926441 | <0.00001 |
| Dvd-CC | -0.50750 | -1.0307356 | 0.01573559 | 0.05882 |

**Table 2.S14. Dunnett test comparison of Cell growth by treatment compared to controls (CC).** Di = Dinn-1, DiD = Dinn-1 diluted to S2 Cq, Dv = Dv-1, DvD = Dv-1 diluted to S2 Cq, Dm = S2.

|  | **Df** | **Sum Sq** | **Mean Sq** | **F-value** | **P-value** |
| --- | --- | --- | --- | --- | --- |
| Treatment | 5 | 25.614 | 5.123 | 15.26 | <0.00001 |
| Residuals | 18 | 6.044 | 0.336 |  |  |

**Table 2.S15. ANOVA test results for Fly Homogenate passage 2 experiment, effect of treatment on cell growth.**

|  | **Diff** | **Lwr** | **Upr** | **Adjusted P** |
| --- | --- | --- | --- | --- |
| Di-CC | 1.04625 | -0.08557568 | 2.1780757 | 0.0754 |
| Did-CC | -0.18875 | -1.32057568 | 0.9430757 | 0.9868 |
| Dm-CC | 0.27875 | -0.85307568 | 1.4105757 | 0.9361 |
| Dv-CC | -1.78000 | -2.91182568 | -0.6481743 | 0.0017 |
| Dvd-CC | 1.46250 | 0.33067432 | 2.5943257 | 0.0093 |

**Table 2.S16. Dunnett test comparison of viral genome change by treatment compared to controls (CC).** Di = Dinn-1, DiD = Dinn-1 diluted to S2 Cq, Dv = Dv-1, DvD = Dv-1 diluted to S2 Cq, Dm = S2.

|  | Df | Sum Sq | Mean Sq | F-value | P-value |
| --- | --- | --- | --- | --- | --- |
| Treatment | 5 | 12.455 | 2.4909 | 4.885 | 0.00535 |
| Residuals | 18 | 9.178 | 0.5099 |  |  |

**Table 2.S17. ANOVA test results for Fly Homogenate passage 2 experiment, effect of treatment on virus genome change per cell.**

|  | **Diff** | **Lwr** | **Upr** | **Adjusted P** |
| --- | --- | --- | --- | --- |
| Di-CC | 1.1950 | -0.199721 | 2.589721 | 0.1076 |
| Did-CC | 0.2025 | -1.192221 | 1.597221 | 0.9928 |
| Dm-CC | 1.2450 | -0.149721 | 2.639721 | 0.0893 |
| Dv-CC | 0.1725 | -1.222221 | 1.567221 | 0.9965 |
| Dvd-CC | 1.9700 | 0.575279 | 3.364721 | 0.0046 |

**Table 2.S18. Dunnett test comparison of viral genome change per cell by treatment compared to controls (CC).** Di = Dinn-1, DiD = Dinn-1 diluted to S2 Cq, Dv = Dv-1, DvD = Dv-1 diluted to S2 Cq, Dm = S2.

|  | **Cell/ Volumes** | **Log_2_ Cells/ Volume** | **Virus/**  **Cell** | **Log_2_ Virus/ Cell** | **Virus/ Volume** | **Log_2_ Virus/ Volume** |
| --- | --- | --- | --- | --- | --- | --- |
| Dinn-1 Control | 6.84 | 2.76 | 0.85 | -0.27 | 5.82 | 2.50 |
| Dinn-1 DiNV | 3.72 | 1.86 | 233.38 | 6.07 | 839.41 | 8.83 |
| Dv-1 Control | 10.56 | 3.40 | 0.06 | -4.19 | 0.58 | -0.80 |
| Dv-1 DiNV | 9.87 | 3.27 | 319.22 | 8.24 | 3145.47 | 11.51 |
| S2 Control | 14.76 | 3.88 | 0.24 | -3.98 | 0.96 | 1.60 |
| S2 DiNV | 24.29 | 4.52 | 0.07 | -2.38 | 5.38 | -0.10 |

**Table 2.S19. Cell-culture adapted virus passage 1 experiment.** Results are reported as either ΔΔCq (log_2_ Cells/ Volume) or 2^ΔΔCq^ (Cells/Volume). Inoculum source is from cell culture adapted virus.

|  | **Df** | **Sum Sq** | **Mean Sq** | **F-value** | **P-value** |
| --- | --- | --- | --- | --- | --- |
| Cells | 2 | 14.254 | 7.127 | 58.257 | <0.00001 |
| Treatment | 1 | 0.103 | 0.103 | 0.840 | 0.37165 |
| Cells:Treatment | 2 | 2.360 | 1.180 | 9.646 | 0.00142 |
| Residuals | 18 | 2.202 | 0.122 |  |  |

**Table 2.S20. Cell-adapted virus passage 1 experiment for cell growth per volume, full model ANOVA results.**

|  | **Diff** | **Lwr** | **Upr** | **Adjusted P** |
| --- | --- | --- | --- | --- |
| Dm-Di | 1.885625 | 1.4392934 | 2.3319566 | <0.00001 |
| Dv-Di | 1.020000 | 0.5736684 | 1.4663316 | 4.53e-05 |
| Dv-Dm | -0.865625 | -1.3119566 | -0.4192934 | <0.00001 |

**Table 2.S21. Tukey test results for cell type from cell-adapted virus passage 1 experiment for host cell increase per volume.**

| **Cell Type** | **Factor** | **Df** | **Sum Sq** | **Mean Sq** | **F-value** | **P-value** |
| --- | --- | --- | --- | --- | --- | --- |
| Dinn-1 | Treatment | 1 | 1.625 | 1.625 | 18.89 | 0.00484 |
|  | Residuals | 6 | 0.516 | 0.086 |  |  |
| Dv-1 | Treatment | 1 | 0.0319 | 0.03188 | 0.402 | 0.55 |
|  | Residuals | 6 | 0.4760 | 0.07933 |  |  |
| S2 | Treatment | 1 | 0.8064 | 0.8064 | 3.999 | 0.0925 |
|  | Residuals | 6 | 1.2101 | 0.2017 |  |  |

**Table 2.S22. Individual ANOVA testing for a treatment effect of viral inoculation on cell increase per volume with each cell type treated separately.**

|  | **Df** | **Sum Sq** | **Mean Sq** | **F-value** | **P-value** |
| --- | --- | --- | --- | --- | --- |
| Cells | 2 | 107.93 | 53.96 | 45.56 | <0.00001 |
| Treatment | 1 | 290.23 | 290.23 | 245.06 | <0.00001 |
| Cells:Treatment | 2 | 102.78 | 51.39 | 43.39 | <0.00001 |
| Residuals | 18 | 21.32 | 1.18 |  |  |

**Table 2.S23. Cell-adapted virus passage 1 experiment for viral genome copy change per volume, full model ANOVA results.**

|  | **Diff** | **Lwr** | **Upr** | **Adjusted P** |
| --- | --- | --- | --- | --- |
| Dm-Di | -4.64375 | -6.032461 | -3.255039 | <0.00001 |
| Dv-Di | -0.30625 | -1.694961 | 1.082461 | 0.84126 |
| Dv-Dm | 4.33750 | 2.948789 | 5.726211 | <0.00001 |

Ta**ble 2.S24. Tukey test results for cell type from cell-adapted virus passage 1 experiment for viral genome increase per volume.**

| **Cell Type** | **Factor** | **Df** | **Sum Sq** | **Mean Sq** | **F-value** | **P-value** |
| --- | --- | --- | --- | --- | --- | --- |
| Dinn-1 | Treatment | 1 | 104.6 | 104.58 | 43.87 | 0.00057 |
|  | Residuals | 6 | 14.3 | 2.38 |  |  |
| Dv-1 | Treatment | 1 | 302.95 | 302.95 | 1119 | <0.00001 |
|  | Residuals | 6 | 1.62 | 0.27 |  |  |
| S2 | 7 | 1 | 9.924 | 9.924 | 13.9 | 0.00976 |
|  | Residuals | 6 | 4.283 | 0.714 |  |  |

**Table 2.S25. Individual ANOVA testing for a treatment effect of viral inoculation on viral increase per volume with each cell type treated separately.**

|  | **Df** | **Sum Sq** | **Mean Sq** | **F-value** | **P-value** |
| --- | --- | --- | --- | --- | --- |
| Cells | 2 | 190.57 | 95.29 | 70.97 | <0.00001 |
| Treatment | 1 | 301.25 | 301.25 | 224.37 | <0.00001 |
| Cells:Treatment | 2 | 117.60 | 58.80 | 43.79 | <0.00001 |
| Residuals | 18 | 24.17 | 1.34 |  |  |

**Table 2.S26. Cell-adapted virus passage 1 experiment for viral genome increase per cell, full model ANOVA results.**

|  | **Diff** | **Lwr** | **Upr** | **Adjusted P** |
| --- | --- | --- | --- | --- |
| Dm-Di | -6.529375 | -8.008016 | -5.0507339 | <0.00001 |
| Dv-Di | -1.326250 | -2.804891 | 0.1523911 | 0.0832139 |
| Dv-Dm | 5.203125 | 3.724484 | 6.6817661 | <0.00001 |

**Table 2.S27. Tukey test results for cell type from cell-adapted virus passage 1 experiment for viral genome increase per cell.**

| **Cell Type** | **Factor** | **Df** | **Sum Sq** | **Mean Sq** | **F-value** | **P-value** |
| --- | --- | --- | --- | --- | --- | --- |
| Dinn-1 | Treatment | 1 | 104.6 | 104.58 | 43.87 | 0.00057 |
|  | Residuals | 6 | 14.3 | 2.38 |  |  |
| Dv-1 | Treatment | 1 | 309.20 | 309.2 | 1562 | <0.00001 |
|  | Residuals | 6 | 1.19 | 0.2 |  |  |
| S2 | Treatment | 1 | 5.072 | 5.072 | 3.508 | 0.11 |
|  | Residuals | 6 | 8.676 | 1.446 |  |  |

**Table 2.S28. Individual ANOVA testing for a treatment effect of viral inoculation on viral genome change per cell with each cell type treated separately.**

| **Cell type** | **Inoculum Source** | **Cell/ Volumes** | **Log_2_ Cells/ Volume** | **Virus/**  **Cell** | **Log_2_ Virus/ Cell** | **Virus/ Volume** | **Log_2_ Virus/ Volume** |
| --- | --- | --- | --- | --- | --- | --- | --- |
| Dinn-1 | Control | 4.645 | 2.215 | 0.344 | -1.540 | 1.602 | 0.675 |
|  | Dinn-1 | 11.151 | 3.146 | 2621.959 | 11.344 | 31567.49 | 14.490 |
|  | Dv-1 | 3.550 | 1.820 | 464.744 | 8.845 | 1631.837 | 10.665 |
|  | S2 | 4.692 | 2.174 | 14.409 | 1.230 | 90.354 | 3.403 |
|  | Diluted Dinn-1 | 5.327 | 2.400 | 201.781 | 7.201 | 1122.722 | 9.601 |
|  | Diluted Dv-1 | 7.616 | 2.904 | 3.284 | -0.644 | 19.170 | 2.260 |
| Dv-1 | Control | 11.374 | 3.506 | 0.175 | -2.543 | 2.000 | 0.964 |
|  | Dinn-1 | 8.262 | 3.025 | 78.331 | 6.288 | 645.420 | 9.313 |
|  | Dv-1 | 9.619 | 3.255 | 186.912 | 7.500 | 1853.557 | 10.755 |
|  | S2 | 8.263 | 3.030 | 1.803 | -1.136 | 12.275 | 1.894 |
|  | Diluted Dinn-1 | 11.034 | 3.446 | 25.995 | 2.681 | 247.418 | 6.128 |
|  | Diluted Dv-1 | 14.642 | 3.840 | 0.555 | -2.640 | 7.306 | 1.200 |
| S2 | Control | 8.938 | 3.123 | 0.152 | -2.784 | 1.289 | 0.339 |
|  | Dinn-1 | 6.535 | 2.673 | 3.357 | 1.713 | 22.630 | 4.385 |
|  | Dv-1 | 5.561 | 2.464 | 0.360 | -1.744 | 1.910 | 0.720 |
|  | S2 | 5.063 | 1.926 | 10.288 | -0.196 | 9.552 | 1.730 |
|  | Diluted Dinn-1 | 7.597 | 2.914 | 0.143 | -3.066 | 1.129 | -0.153 |
|  | Diluted Dv-1 | 14.528 | 3.839 | 0.040 | -4.671 | 0.564 | -0.833 |

**Table 2.S29. Cell-culture adapted virus passage 2 experiment in Dinn-1 cells.** Results are reported as either ΔΔCq (e.g. Log_2_ Cells/ Volume) or 2^ΔΔCq^ (e.g. Cells/Volume). Inoculum source is from cell culture adapted phase 1 experiment.

| **Metric** | **Factor** | **Df** | **Sum Sq** | **Mean Sq** | **F-value** | **p-value** |
| --- | --- | --- | --- | --- | --- | --- |
| Cells per volume | Treatment | 5 | 4.885 | 0.9770 | 3.297 | 0.0275 |
|  | Residuals | 18 | 5.334 | 0.2963 |  |  |
| Virus per volume | Treatment | 5 | 606.3 | 121.25 | 27.79 | <0.00001 |
|  | Residuals | 18 | 78.6 | 4.36 |  |  |
| Virus per cell | Treatment | 5 | 586.4 | 117.27 | 28.76 | <0.00001 |
|  | Residuals | 18 | 73.4 | 4.08 |  |  |

**Table 2.S30. ANOVA results for cell-adapted virus study passage 2 in Dinn-1 cells for each metric separately.**

| **Metric** | **Comparison** | **Diff** | **Lower** | **Upper** | **p-value** |
| --- | --- | --- | --- | --- | --- |
| Cells per volume | Di-CC | 0.93125 | -0.1319702 | 1.9944702 | 0.0973 |
|  | Did-CC | 0.18500 | -0.8782202 | 1.2482202 | 0.9842 |
|  | Dm-CC | -0.04125 | -1.1044702 | 1.0219702 | 1.0000 |
|  | Dv-CC | -0.39500 | -1.4582202 | 0.6682202 | 0.7578 |
|  | Dvd-CC | 0.68875 | -0.3744702 | 1.7519702 | 0.2929 |
| Virus per volume | Di-CC | 13.81500 | 9.734793 | 17.895207 | <0.00001 |
|  | Did-CC | 8.92625 | 4.846043 | 13.006457 | 0.00004 |
|  | Dm-CC | 2.72875 | -1.351457 | 6.808957 | 0.2673 |
|  | Dv-CC | 9.99000 | 5.909793 | 14.070207 | 0.00005 |
|  | Dvd-CC | 1.58500 | -2.495207 | 5.665207 | 0.7273 |
| Virus per cell | Di-CC | 12.88375 | 8.939478 | 16.828022 | <0.00001 |
|  | Did-CC | 8.74125 | 4.796978 | 12.685522 | <0.00001 |
|  | Dm-CC | 2.77000 | -1.174272 | 6.714272 | 0.2296 |
|  | Dv-CC | 10.38500 | 6.440728 | 14.329272 | <0.00001 |
|  | Dvd-CC | 0.89625 | -3.048022 | 4.840522 | 0.9529 |

**Table 2.S31. Dunnett’s test results for cell-adapted virus study passage 2 in Dinn-1 cells for each metric separately.**

| **Metric** | **Factor** | **Df** | **Sum Sq** | **Mean Sq** | **F-value** | **p-value** |
| --- | --- | --- | --- | --- | --- | --- |
| Cells per volume | Treatment | 5 | 1.963 | 0.3927 | 6.101 | 0.00179 |
|  | Residuals | 18 | 1.158 | 0.0644 |  |  |
| Virus per volume | Treatment | 5 | 373.4 | 74.69 | 20.02 | <0.00001 |
|  | Residuals | 18 | 67.1 | 3.73 |  |  |
| Virus per cell | Treatment | 5 | 402.1 | 80.42 | 19.12 | <0.00001 |
|  | Residuals | 18 | 75.7 | 4.21 |  |  |

**Table 2.S32. ANOVA results for cell-adapted virus study passage 2 in Dv-1 cells for each metric separately.**

| **Metric** | **Comparison** | **Diff** | **Lower** | **Upper** | **p-value** |
| --- | --- | --- | --- | --- | --- |
| Cells per volume | Di-CC | -0.48125 | -0.976746 | 0.01424599 | 0.0585 |
|  | Did-CC | -0.06000 | -0.555496 | 0.43549599 | 0.9969 |
|  | Dm-CC | -0.47625 | -0.971746 | 0.01924599 | 0.0618 |
|  | Dv-CC | -0.25125 | -0.746746 | 0.24424599 | 0.5087 |
|  | Dvd-CC | 0.33375 | -0.161746 | 0.82924599 | 0.2612 |
| Virus per volume | Di-CC | 8.34875 | 4.576762 | 12.120738 | 0.00003 |
|  | Did-CC | 5.16375 | 1.391762 | 8.935738 | 0.0059 |
|  | Dm-CC | 0.93000 | -2.841988 | 4.701988 | 0.9358 |
|  | Dv-CC | 9.79125 | 6.019262 | 13.563238 | <0.00001 |
|  | Dvd-CC | 0.23625 | -3.535738 | 4.008238 | 0.9999 |
| Virus per cell | Di-CC | 8.83000 | 4.824319 | 12.835681 | 0.00005 |
|  | Did-CC | 5.22375 | 1.218069 | 9.229431 | 0.0087 |
|  | Dm-CC | 1.40625 | -2.599431 | 5.411931 | 0.7932 |
|  | Dv-CC | 10.04250 | 6.036819 | 14.048181 | 0.00001 |
|  | Dvd-CC | -0.09750 | -4.103181 | 3.908181 | 1.0000 |

**Table 2.S33. Dunnett’s test results for cell-adapted virus study passage 2 in Dv-1 cells for each metric separately.**

| **Metric** | **Factor** | **Df** | **Sum Sq** | **Mean Sq** | **F-value** | **p-value** |
| --- | --- | --- | --- | --- | --- | --- |
| Cells per volume | Treatment | 5 | 8.342 | 1.6684 | 3.679 | 0.0181 |
|  | Residuals | 18 | 8.163 | 0.4535 |  |  |
| Virus per volume | Treatment | 5 | 68.75 | 13.750 | 10.78 | 0.00006 |
|  | Residuals | 18 | 22.96 | 1.276 |  |  |
| Virus per cell | Treatment | 5 | 102.91 | 20.582 | 7.65 | 0.00052 |
|  | Residuals | 18 | 48.43 | 2.691 |  |  |

**Table 2.S34. ANOVA results for cell-adapted virus study passage 2 in S2 cells for each metric separately.**

| **Metric** | **Comparison** | **Diff** | **Lower** | **Upper** | **p-value** |
| --- | --- | --- | --- | --- | --- |
| Cells per volume | Di-CC | -0.45000 | -1.7653365 | 0.8653365 | 0.8081 |
|  | Did-CC | -0.20875 | -1.5240865 | 1.1065865 | 0.9894 |
|  | Dm-CC | -1.19625 | -2.5115865 | 0.1190865 | 0.0819 |
|  | Dv-CC | -0.65875 | -1.9740865 | 0.6565865 | 0.5198 |
|  | Dvd-CC | 0.71625 | -0.5990865 | 2.0315865 | 0.4444 |
| Virus per volume | Di-CC | 4.04625 | 1.8402155 | 6.252285 | 0.0004 |
|  | Did-CC | -0.49125 | -2.6972845 | 1.714785 | 0.9564 |
|  | Dm-CC | 1.39125 | -0.8147845 | 3.597285 | 0.3151 |
|  | Dv-CC | 0.38125 | -1.8247845 | 2.587285 | 0.9846 |
|  | Dvd-CC | -1.17125 | -3.3772845 | 1.034785 | 0.46710 |
| Virus per cell | Di-CC | 4.49625 | 1.2925075 | 7.699992 | 0.0047 |
|  | Did-CC | -0.28250 | -3.4862425 | 2.921242 | 0.9993 |
|  | Dm-CC | 2.58750 | -0.6162425 | 5.791242 | 0.1385 |
|  | Dv-CC | 1.04000 | -2.1637425 | 4.243742 | 0.8362 |
|  | Dvd-CC | -1.88750 | -5.0912425 | 1.316242 | 0.3738 |

**Table 2.S35. Dunnett’s test results for cell-adapted virus study passage 2 in S2 cells for each metric separately.**

### Supplemental Figures

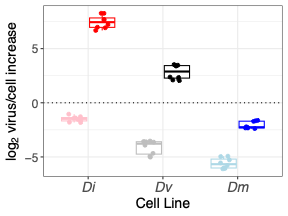

**Figure S1.** **Passage 1 with fly homogenate virus log_2_ genome fold increase per cell (ΔΔCq).** Cell types are: Dinn-1 (Di, *D. innubila*), Dv-1 (Dv, *D. virilis*) and S2 (Dm, *D. melanogaster*). Lighter colors are mock infected and darker colors are infected with virus.

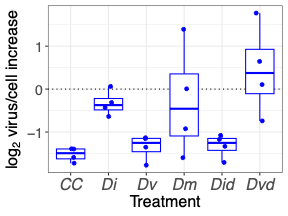

**Figure S2. Log_2_ virus per cell genome fold increase from fly homogenate passage 2 experiment.** Cell type is Dinn-1. Treatment is passage 1 virus derived from the passage 1 study (mock = CC, Di from *D. innubila*, Dv from *D.* *virilis*, Dm from *D. melanogaster*, Did from diluted *D. innubila*, Dvd from diluted *D. virilis.*

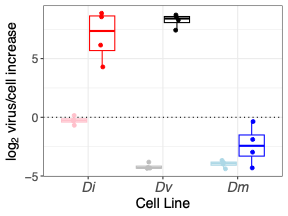

Figure S3. Cell Culture Adapted passage 1 Study log_2_ virus per cell genome increase. Cell types are: Dinn-1 (Di, D. innubila), Dv-1 (Dv, D. virilis) and S2 (Dm, D. melanogaster). Lighter colors represent mock inoculated and darker colors represent DiNV inoculated.

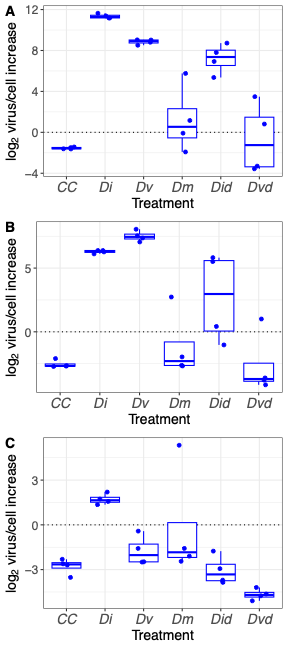

**Figure S4. Cell-culture adapted passage 2 results for log_2_ virus per cell**. A) Dinn-1 cells, B) Dv-1 cells, C) S2 cells. Treatment is passage 1 virus derived from the cell-culture adapted passage 1 study (mock = CC, Di from *D. innubila*, Dv from *D. virili*s, Dm from *D. melanogaster*, Did from diluted *D. innubila*, Dvd from diluted *D. virilis*).

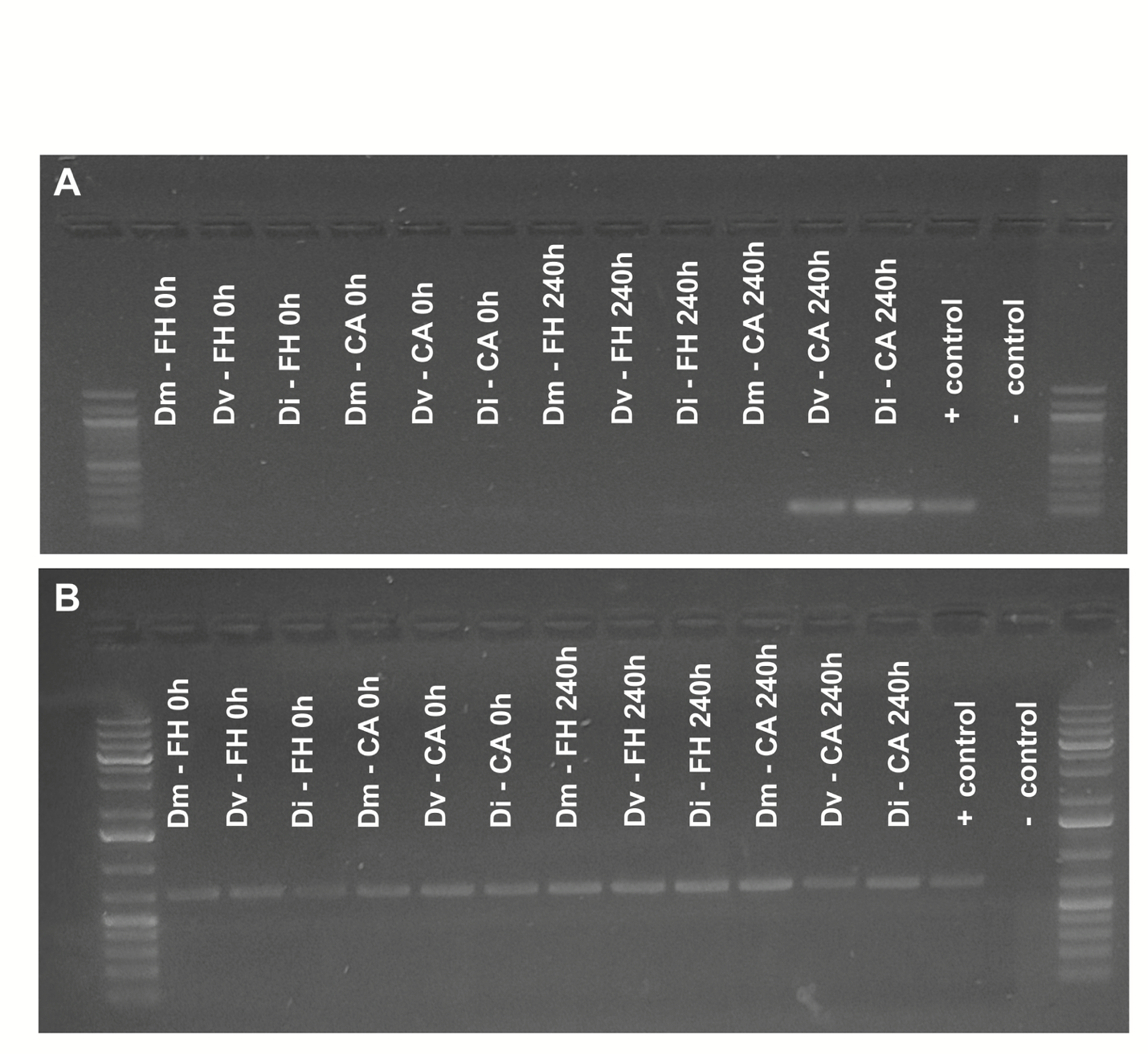

**Figure S5. Viral transcripts only evident at 240-hours post inoculation in adapted virus for *D. innubila* and *D. virilis* samples**. RT-PCR results for A) viral *PIF3* gene and B) host *COI* at 0 or 240 hours post inoculation with naïve (fly homogenate, FH) or adapted (cell adapted, CA) virus in *D. innubila* (Di), *D. virilis* (Dv) or *D. melanogaster* (Dm) cells. Ladders are on either ends (100bp in A and 1kb+ in B) and + control refers to a known infected sample and negative control is water.

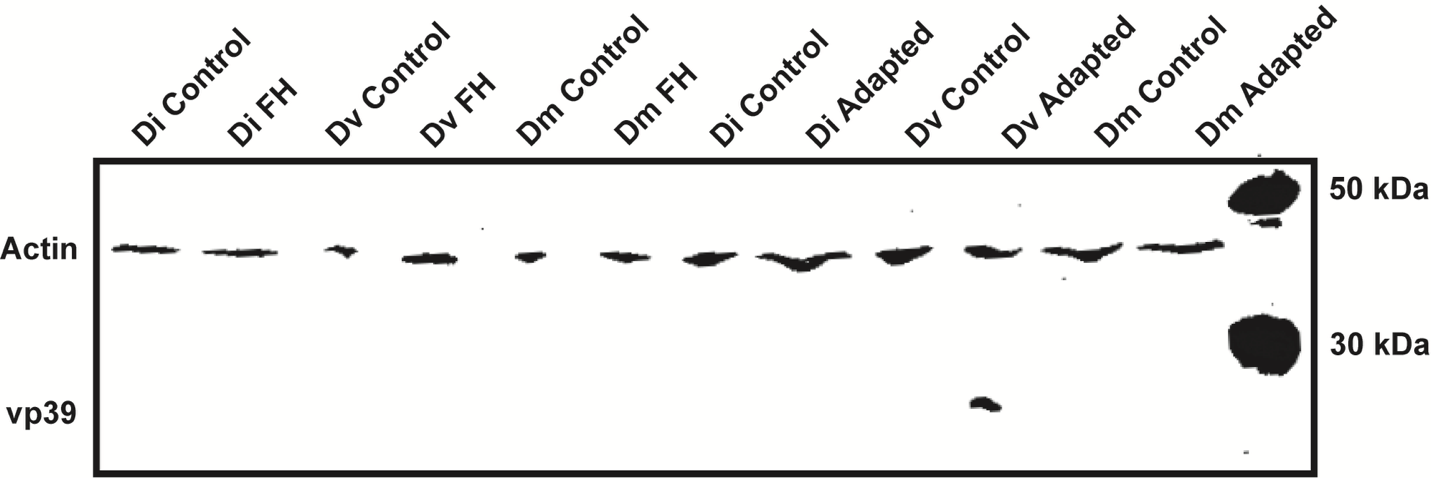

**Figure S6. Viral capsid protein is only evident at 240-hours post inoculation in adapted virus for *D. D. virilis* samples**. Western blot of anti-vp39 and anti-actin antibodies at 240 hours post inoculation with naïve (fly homogenate, FH) or adapted (cell adapted, CA) virus in *D. innubila* (Di), *D. virilis* (Dv) or *D. melanogaster* (Dm) cells or in controls (only cell culture media added).
